# Raman-probe-derived biomarkers account for cartilage composition, structure, and viscoelastic properties

**DOI:** 10.64898/2026.10.30.762336

**Authors:** Massie Kazemi, Juuso Tuppurainen, Jiri Jäntti, Chenhao Yu, Maria C. Fugazzola, René V. Weeren, Thomas P. Schaer, Mark W. Grinstaff, Mads S. Bergholt, Brian D. Snyder, Janne T.A. Mäkelä, Michael B. Albro

## Abstract

Raman spectroscopy is an inelastic light scattering technique that reflects the vibrational modes of the biochemical building blocks of the major constituents that comprise the cartilage extracellular matrix (ECM)—glycosaminoglycans (GAG), type-II collagen (COL), and water. We investigated whether ECM-specific biomarkers measured with our recently developed arthroscopy-compatible Raman needle probe characterize the functional viscoelastic properties of articular cartilage explants. Equine osteochondral explants were harvested from high-load (femoral condyle) and low-load (intertrochlear groove) sites to evaluate Raman-derived biomarkers of cartilage structure and function. Viscoelastic material properties (equilibrium/dynamic modulus, phase shift) were determined by mechanical indentation. A computational fibril-reinforced biphasic material model was used to estimate the glycosaminoglycan-associated non-fibrillar-matrix modulus, the collagen-associated fibril-network modulus, and hydraulic permeability. Raman-derived biomarkers enabled evaluation of cartilage tissue composition and structure, accounting for 81% of the variation in GAG content, 53% of the variation in COL content, and 58% of the variation in COL anisotropy. Biomarkers also accounted for 81% of the variation in equilibrium modulus, 58% in dynamic modulus, and 51% in phase shift. For poroelastic properties, biomarkers accounted for 83% of the variation in non-fibrillar-matrix modulus, 25% of the variation in fibril-network modulus, and 34% of the variation in hydraulic permeability.

**Clinical Significance:** Raman spectroscopy provides objective compositional biomarkers that account for viscoelastic material properties underlying articular cartilage function. These findings support its future use as an arthroscopic platform for objective assessment of cartilage injury, monitoring of disease progression, and evaluation of therapeutic interventions.

## Introduction

Hyaline cartilage is the hydrated, avascular connective tissue that provides a smooth, nearly frictionless surface while supporting applied compressive loads during diarthrodial joint motion. The cartilage extracellular matrix (ECM) is comprised of an anisotropic type-II collagen fibril (COL) network (5-20% per wet weight) that affords structure and tensile strength, complemented by a negatively charged, sulfated glycosaminoglycan (GAG) matrix (5-10% per wet weight) that retains high levels of interstitial water (70-90% per wet weight).^1^ These components act synergistically, imparting the viscoelastic and tribological material properties essential to cartilage function. The fixed negative charges of the GAG matrix impart a Donnan osmotic swelling pressure ^2;3^ that is restrained by the tensile collagen fibrils. In response to compressive joint loads, the collagen network deforms, decreasing pore volumes, leading to an increase in the local anionic fixed charge density that acts to slow interstitial fluid transport and lower hydraulic tissue permeability. Pressurization of the entrapped fluid provides interstitial fluid load support (IFLS), supporting >90% of compressive joint forces ^4;5^. This strain-dependent decrease in tissue permeability and the increased viscous drag of interstitial fluid flowing through the compressed porous collagen network account for the tissue’s non-linear, viscoelastic behavior that serves to dampen tissue strains and dissipate energy.^6;7^ Molecular rearrangement of the deformed solid matrix affords additional flow-independent viscoelasticity.^8;9^ Through these mechanistic structure-function relationships, GAG and collagen contents generally account for 30% to 70% of the cartilage’s functional viscoelastic properties.^10–12^

Focal articular cartilage lesions are common musculoskeletal injuries resulting from joint trauma, such as ligament rupture, meniscal tears, or chondral contusions.^13^ Mechanical and chemical insults induce progressive compositional changes, including loss of GAG, disruption of the collagen network architecture, and tissue swelling, collectively leading to surface fibrillation and structural failure at lesion sites. ^14;15^ The loss of anionic GAGs and collagen network degradation diminish Donnan swelling pressure and increase interstitial fluid permeability, thereby reducing IFLS ^16;17^ and the viscous drag of interstitial fluid flowing through the porous collagen network. These compositional and structural changes degrade both the viscoelastic and equilibrium material properties critical to load-bearing joint function.^18^ Without intervention, cartilage lesions progress to post-traumatic osteoarthritis, making early repair essential. Current surgical strategies (e.g., microfracture and osteochondral autograft transfer) aim to restore joint function, yet their success remains variable and difficult to predict.

Successful cartilage repair requires objective intraoperative assessment of lesion composition and mechanical competency. Current diagnostic approaches have significant limitations. Arthroscopic cartilage grading schemes (e.g., Outerbridge) rely on subjective visual assessment of surface topology and poorly correlate with tissue function.^19^ Advanced imaging modalities (e.g., compositional MRI (T1ρ, T2, T2* mapping),^20;21^, ultrasound,^22^ and optical coherence tomography (OCT)^23^) provide only moderate correlation with cartilage composition and material behavior, and have limited intraoperative applicability. Histopathological grading schemes (e.g., OARSI, Mankin) accurately portray the pathoanatomic stages of OA,^24^ but require invasive tissue sampling and tissue destruction, making it unsuitable for clinical assessments of cartilage competency. Thus, there exists a critical unmet need for non-destructive, interoperative techniques to objectively quantify cartilage composition while accounting for functional biomechanical properties.

Vibrational spectroscopy techniques offer promise for non-destructive compositional assessment during arthroscopic procedures.^25–34^ Raman spectroscopy is predicated on the inelastic photon scattering induced by monochromatic laser irradiation, which produces wavelength shifts corresponding to specific molecular vibrational modes.^35^ The resulting Raman spectrum provides a quantitative optical fingerprint of tissue molecular composition (e.g., amides, sulfates, carboxylic acids, and hydroxyls) that directly identifies biochemical constituents (GAG, COL, water) of the cartilage ECM ^36–39^ We previously developed an arthroscopy-compatible Raman spectroscopic needle probe that enables a real-time, non-destructive optical biopsy of cartilage during arthroscopic procedures.^40;41^ In our prior studies, Raman-probe-derived compositional biomarkers accounted for 70%-80% of the variation in cartilage GAG and water content.^42^ This platform had been used to assess compositional changes during *in vitro* cultivation of tissue engineered cartilage ^43^ and *in vivo* site-specific evaluations of cartilage composition in pre-clinical animal models after treatment by microfracture or autologous cell implantation.^44^ Despite these advances, we have yet to determine whether Raman-probe-derived compositional biomarkers portray the viscoelastic and load-bearing functional properties of articular cartilage.

To address this question, we investigated whether Raman-probe-derived compositional biomarkers for GAG, COL, and water account for the viscoelastic functional properties (equilibrium elastic modulus, dynamic modulus, phase shift) of articular cartilage. The analysis utilized cartilage explants from high load bearing (medial femoral condyle) and low load bearing (intertrochlear groove) anatomic regions in the equine stifle joint ^45;46^. While these tissues do not possess macroscopic cartilage lesions, they possess a range of compositions consistent with changes that occur in early-stage cartilage degeneration, thereby enabling validation of Raman sensitivity to describe subtle but physiologically relevant variations in cartilage composition, structure, and function. Previously, this model has demonstrated a strong relationship between tissue composition and viscoelastic tissue properties.^46^ Structure and composition were analyzed on tissue sections using digital densitometry, Fourier-transform infrared spectroscopy, and polarized light microscopy. Viscoelastic material properties were determined from indentation stress relaxation tests. Further, biphasic fluid–solid interactions were modeled using fibril-reinforced poroelastic finite element analysis to quantify the proteoglycan-associated non-fibrillar matrix modulus, the collagen-associated fibril network modulus, and interstitial fluid permeability.

## Methods

### Tissue samples

Twelve equine stifle joints from 4-6-year-old livestock Criollo horses (from abattoir) were selected, excluding those with visible abnormalities. For each animal, either the left or the right stifle joint was used. Information regarding the sex of the animals was not available. Osteochondral plugs, 8.5 mm in diameter, were extracted using a cylindrical biopsy saw (**Fig. 1**) from the load bearing (LB) medial femoral condyle (n = 12) and the non-load bearing (NLB) distal intertrochlear groove of the femoropatellar joint (n = 12).^45–47^ The chondral layer thickness of plugs was evaluated via μCT imaging, exhibiting values of 1.70 ± 0.47 mm and 1.33 ± 0.30 mm for femoral condyle and intertrochlear groove sites, respectively.^45^

**Figure 1:**
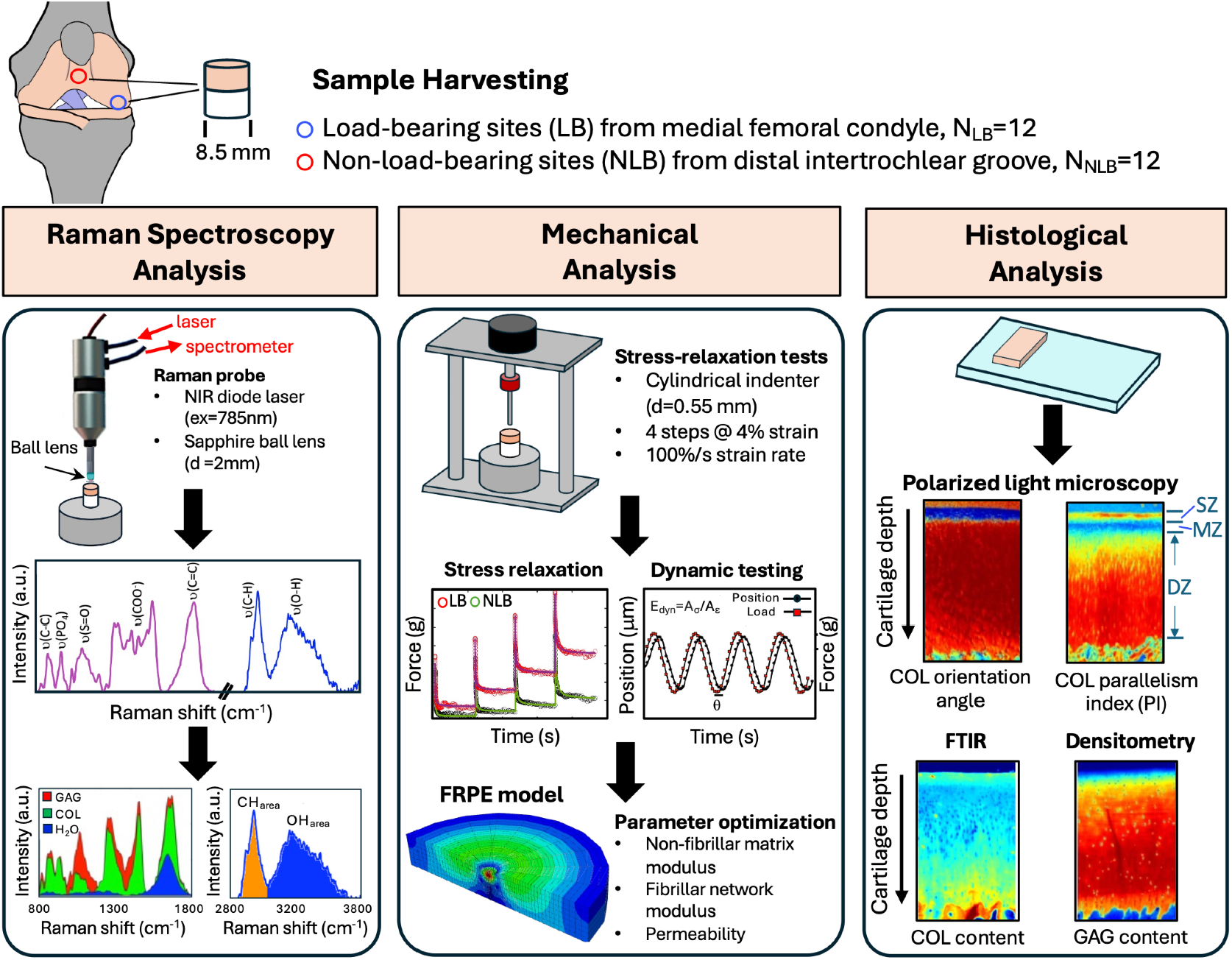
Experimental workflow. Protocols for equine osteochondral explant analysis.

### Raman analysis

The Raman platform consists of a custom needle probe that transmits an excitation NIR diode laser (ex=785nm, 500mW, B&WTek) through a Ø2mm sapphire ball lens. Raman scattered light reflected from the cartilage is passed through the ball lens via a dichroic mirror, across a long pass filter coupled to a spectrometer (Eagle Raman-S, Ibsen Photonics) equipped with a NIR optimized charge coupled device (Ivac, Andor Oxford Instruments). High-quality spectral measurements were acquired with the Ø2mm ball lens in gentle contact with the tissue surface over a 30-second integration time corresponding to an approximate laser footprint area of 0.30 mm^2^, with ~300 μm depth of penetration into the tissue.^40–42^ The spectral fingerprint range (800-1800cm^−1^) is preprocessed using a 3^rd^ order polynomial baseline subtraction, followed by area-under-curve normalization. The resulting cartilage spectral profile **Cartilage**_spectra_ is decomposed via a reference-based multivariate linear regression model:

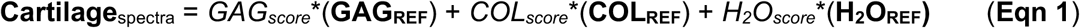

where the derived regression coefficients (*GAG*_*score*_, *COL*_*score*_, *H*_*2*_*O*_*score*_) represent the relative contribution of the predominant constituents of cartilage ECM (GAG, COL, H_2_O) to the composite cartilage Raman spectra (**Fig. 1**), calculated using the spectra of purified reference chemicals for each ECM constituent: **GAG**_**REF**_=chondroitin sulfate (shark cartilage; Sigma[C4384]); **COL**_**REF**_=type-II collagen (chicken sternal cartilage; Sigma[C9301]); and **H**_**2**_**O**_**REF**_=PBS. For each constituent, the Raman score ranges from a value of 0 to 1 based on the constituent’s weighted contribution to the tissue spectra. Additionally, the high-wavenumber range (2700-3800 cm^−1^) is used to compute the ratio of areas under the OH and CH_2_ peaks, reflecting tissue hydration and organic ECM components, respectively, which were similarly preprocessed using a linear baseline subtraction, area-under-curve normalization, followed by measurement of the area under the organic-content-associated CH_2_ region (CH_area_) and water-content-associated OH region (OH_area_).^27^

### Compositional analysis

Cartilage is organized into distinct zones—superficial, middle, and deep— that vary in depth and collagen orientation: *Superficial Zone*, 10%–20% of the total thickness, with collagen fibrils oriented parallel to the surface; *Middle Zone*, 10%–40% of the total thickness, with more random collagen spacing; and *Deep Zone*, approximately 50-70% of the total thickness, characterized by radially oriented bundles of fibrils that cross the tidemark.^46^ GAG content is also zonally distributed, increasing with depth.^48^ From histological sections, the spatial distributions of GAG and collagen were determined by digital densitometry and Fourier-transform infrared spectroscopy (FTIR), respectively.^45;46^ (**Fig.1**). For digital densitometry, Safranin-O-stained sections were imaged and grayscale values were converted to optical density using calibration filters to enable quantitative assessment of GAG content. The collagen parallelism index, representing material anisotropy, was determined using polarized light microscopy.^45;49^ Collagen content was quantified from FTIR images by integrating the absorbance of the amide I region (1595–1720 cm^−1^), which correlates with collagen content. ^50;51^ For each modality, depth-wise profiles were generated by averaging pixels at the same tissue depth and subsequently averaged across sections. The spatial distributions of GAG and collagen content were averaged over the uppermost 300 μm of the chondral surface, the depth of interrogation by the Raman probe, to facilitate correlative validations.

### Biomechanical property measures

Functional material properties (equilibrium elastic modulus, dynamic modulus, phase shift) were derived using a 4-step stress-relaxation test performed using a flat-end cylindrical indenter (Ø=0.55 mm) (applied perpendicular to the chondral surface, positioned with a goniometer).^46;47^ Screws inserted into the subchondral bone anchored the samples. After immersion in PBS (310 mOsmmol/L), a 40 kPa pre-stress was applied to ensure initial contact between the indenter and chondral surface, resulting in cartilage deformations ranging from 8 to 60µm at equilibrium. The 4-step prescribed strain stress relaxation test consisted of compressing the cartilage by 4% of the cartilage thickness at a ramp rate of 100% per second, followed by a 600-second relaxation period, repeated over four cycles. The equilibrium Young’s modulus (*E*_eq_) was obtained by applying linear least-squares regression to the stress–strain data collected at the equilibrium points of each loading step. After the stress-relaxation test, a sinusoidal dynamic test without lift-off was conducted at the equilibrium indentation depth reached at the end of the final relaxation step, with a 4% peak-to-peak strain amplitude and 4 cycles at 1 Hz. The dynamic modulus *E*_dyn_ was calculated as the ratio of stress-strain amplitude, and the phase shifts (*θ*) were determined using a Fourier transform. The phase shift describes the viscoelastic lag between stress and strain, where larger *θ* values indicate greater viscous energy dissipation relative to elastic energy storage.^52^

A phase shift of *θ* = 0° represents a fully elastic material, whereas *θ* = 90° represents a fully viscous material.

### Fibril-reinforced poroelastic analysis

Five effective material parameters were estimated using inverse finite element optimization by minimizing the mean-squared relative error between experimentally measured and simulated reaction forces. The optimized parameters included non-fibrillar matrix modulus (*E*_nf_), initial fibrillar network modulus (*E*_f_^0^; referred to hereafter as the fibrillar network modulus), strain-dependent fibrillar network modulus (*E*_f_^ε^), permeability strain-dependency factor (*M*), and initial Darcy permeability (*k*_0_; referred to hereafter as Darcy permeability). These are not intrinsic constituent-level properties (e.g., true collagen fibril stiffness), but effective bulk parameters of the fibril-reinforced continuum. Parameter estimation was performed using sample-specific, axisymmetric fibril-reinforced biphasic finite element (FE) model (Abaqus v6.12-3, Dassault Systèmes, Vélizy-Villacoublay, France).^47^ The material model was developed based on the work of Li et al. and Wilson et al.^53;54^ The collagen fibril network was modeled as a nonlinear elastic fibrillar matrix.^52;54;55^ The non-fibrillar matrix was modeled as Neo-Hookean hyperelastic material.^52;53^ Darcy’s law was used to characterize fluid flow within the porous matrix. Consistent with prior FE models, while the thickness of each sample was modeled individually, the internal composition and void ratio were assumed to be homogeneous to determine the material properties independent of spatial heterogeneity.^52;55–57^ The total stress tensor was computed as the combination of stress tensors arising from both the fibrillar and non-fibrillar matrices, as well as the pore fluid pressure.^52;58^

### Statistical analysis

Paired t-tests (two-sided, significance level p < 0.05) were used to compare LB vs. NLB specimens, accounting for nesting of data (one LB and one NLB specimen) within animals. Bivariate regression analyses (linear and polynomial) were used to examine associations between Raman-derived biomarkers and compositional/structural/mechanical properties. When polynomial regression yielded a non-monotonic relationship over the measured range, linear regression was used instead. Statistical analyses were performed using Stata software (v18.1, StataCorp LLC, College Station, Texas).

## Results

### Composition & structure

Explants from LB and NLB sites exhibited differences in tissue composition and structure consistent with the depth dependent variation in zonal anatomy **(Fig. 2)**. Histological analysis of the full chondral layer revealed that NLB trochlear cartilage compared to LB medial condyle cartilage displayed less GAG **(Fig. 2A)**, increased COL **(Fig. 2B)**, and greater superficial zone anisotropy (increased collagen parallelism index) (**Fig. 2C**). Quantitative histological analysis of the uppermost 300 μm of the chondral layer corresponding to the depth of interrogation by the Raman probe was performed to facilitate comparison to Raman spectra derived biomarkers representing GAG, COL and water ECM content. Like the full depth chondral tissue analysis, the superficial and middle zones of NLB cartilage compared to LB cartilage demonstrated significantly less GAG content (**Fig. 2D;** *p* < 0.05), significantly more collagen content (**Fig. 2E**; *p* < 0.05), and significantly more anisotropy (parallelism index) (**Fig. 2F**; *p* < 0.05).

**Figure 2:**
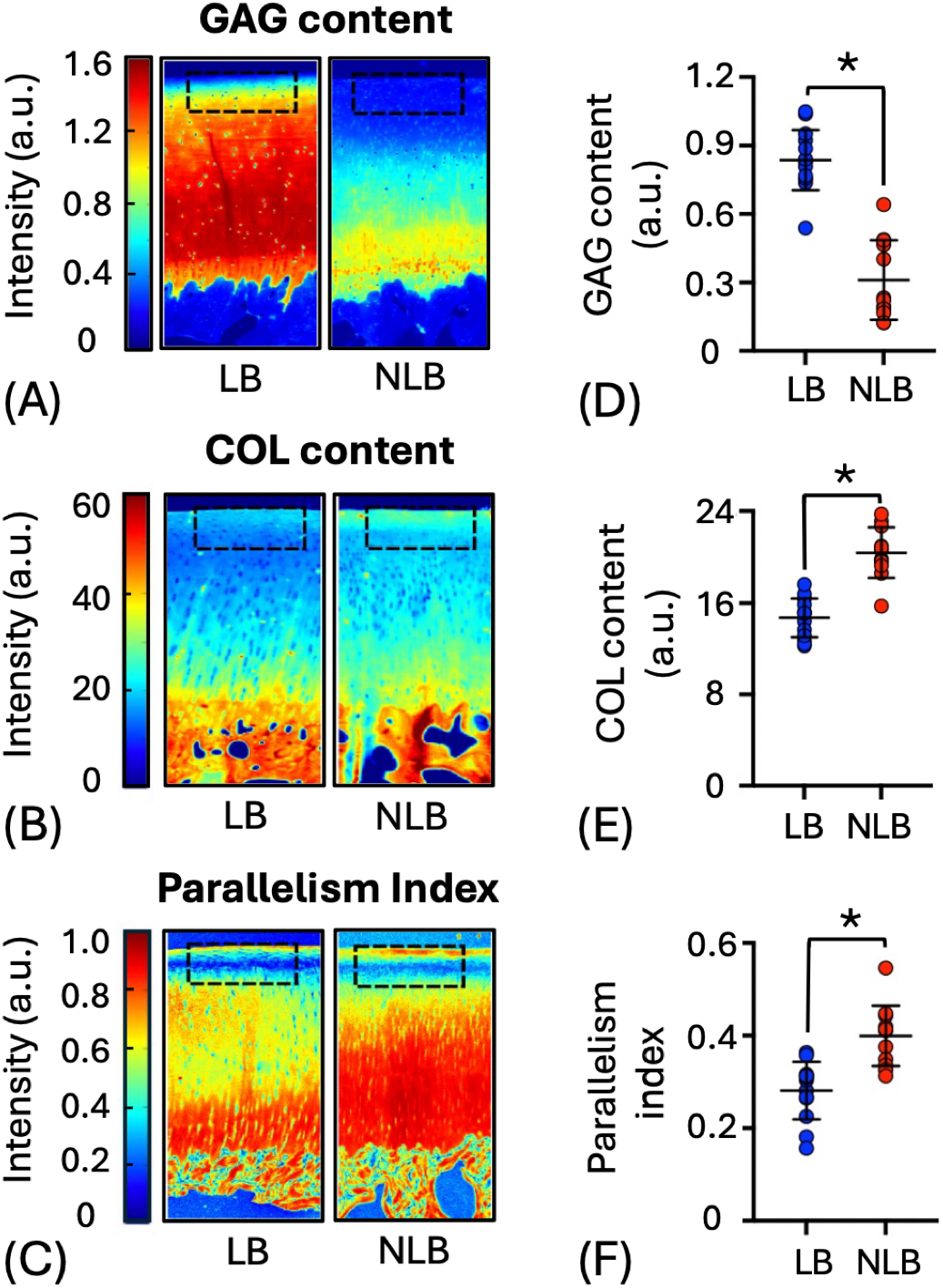
Composition and structure analysis of LB and NLB equine osteochondral explants. Representative images depicting relative distribution of (A) GAG via digital densitometry (B) collagen via FTIR, and (C) collagen parallelism index via polarized light microscopy through full thickness of chondral layer in LB and NLB explants. (D) GAG content, (E) collagen content, and (F) collagen parallelism index in LB and NLB explants within the topmost 300μm of chondral layer (highlighted by black dashed rectangles in A-C). \**p* < 0.05

### Viscoelastic & poroelastic properties

The equilibrium elastic modulus of NLB cartilage was significantly less compared to LB cartilage (**Fig.3A;** *p* < 0.05). Response to dynamic mechanical loading showed a similar trend where the dynamic modulus of NLB cartilage was less than LB cartilage but did not quite reach significance (**Fig.3B;** *p* = 0.06); however, there was a significant increase in the phase shift angle for NLB compared to LB cartilage (**Fig.3C**; *p* < 0.05), indicating more viscous behavior from trochlear groove cartilage. Among the biphasic parameters derived from the FE model, the GAG-association-non-fibrillar matrix modulus was significantly lower for NLB cartilage compared to LB cartilage (**Fig.3D**; *p* < 0.05). There were no significant changes in the fibril network modulus between tissue sites (**Fig.3E**; *p* = 0.21). The Darcy permeability of LB cartilage trended lower than NLB cartilage but did not quite reach significance (**Fig.3F**; *p* = 0.06).

**Figure 3:**
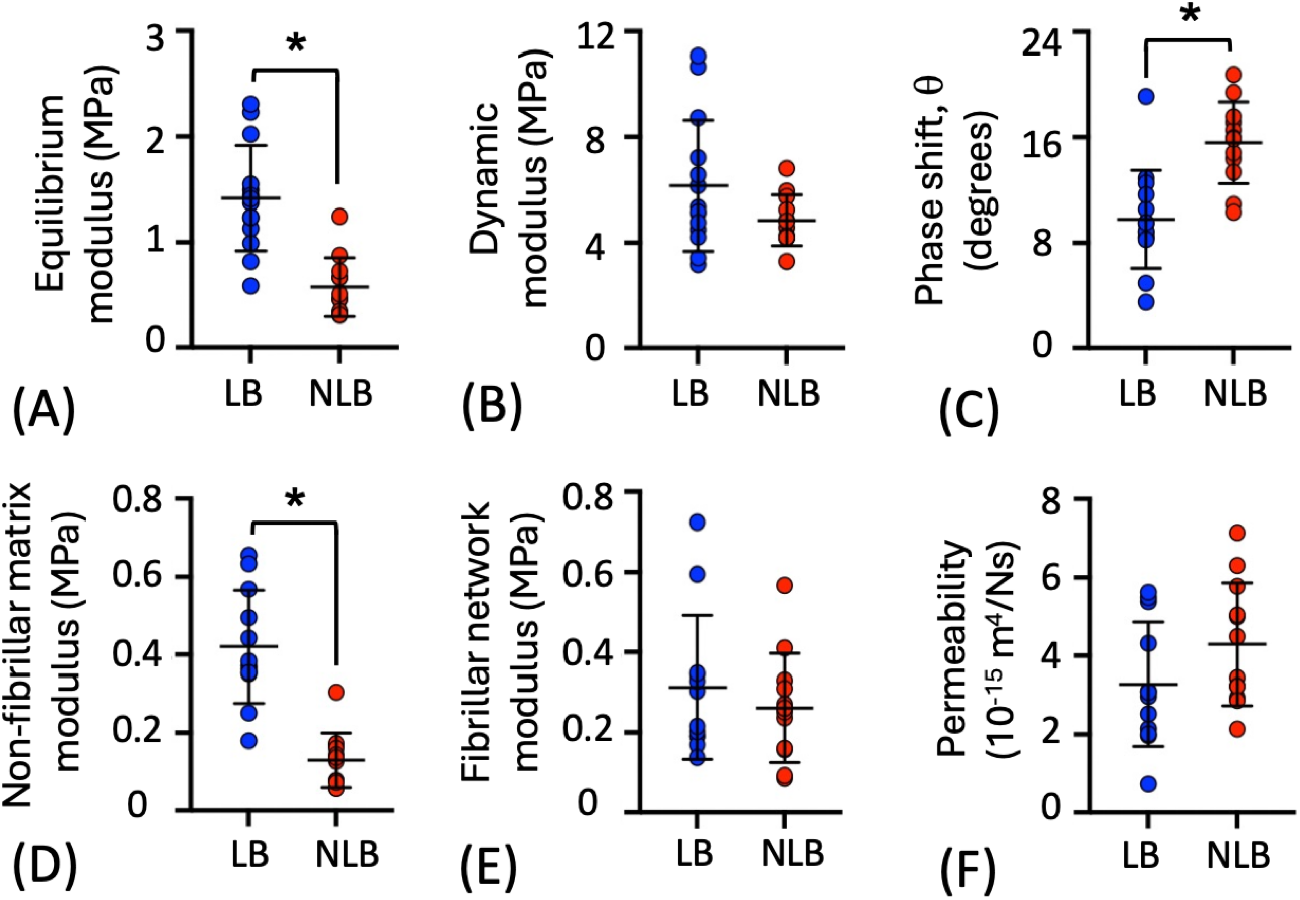
Viscoelastic and biphasic properties of LB and NLB equine osteochondral explants. (A) Equilibrium modulus, (B) dynamic modulus, (C) phase shift, (D) non-fibrillar matrix modulus, (E) fibril network modulus, (F) Darcy permeability. \**p* < 0.05. To accommodate the increased load bearing requirement of the medial condyle cartilage (LB) relative to the trochlear groove cartilage (NLB), the LB cartilage is stiffer and more elastic (lower phase shift angle, p <0.05) due to higher GAG content contributing to the stiffer non-fibrillar matrix modulus, with a tendency to be less permeable (permeability p=0.06) to viscous fluid flow.

### Raman biomarkers

Multivariate decomposition of the composite cartilage spectral profile into the predominant cartilage ECM constituents (GAG, COL, H_2_O) accounted for 92 ± 2% of the variation of the fingerprint Raman spectra. The Raman-derived biomarkers portrayed the same differences in tissue composition and structure between LB medial condyle explants versus NLB trochlea explants, as established by quantitative histological analysis of the superficial and middle zones: GAG_score_ was significantly lower for NLB explants, relative to LB explants (**Fig.4E**; p < 0.05); COL_score_ was significantly higher for NLB explants, relative to LB explants (**Fig.4F**; *p* < 0.05); H_2_O_score_ was significantly lower for NLB explants, relative to LB explants (**Fig.4G**; *p* < 0.05). Analysis of the Raman high wavenumber region revealed no significant difference in OH_area_ and CH_area_ between tissue regions (**Fig.4H&I**).

**Figure 4:**
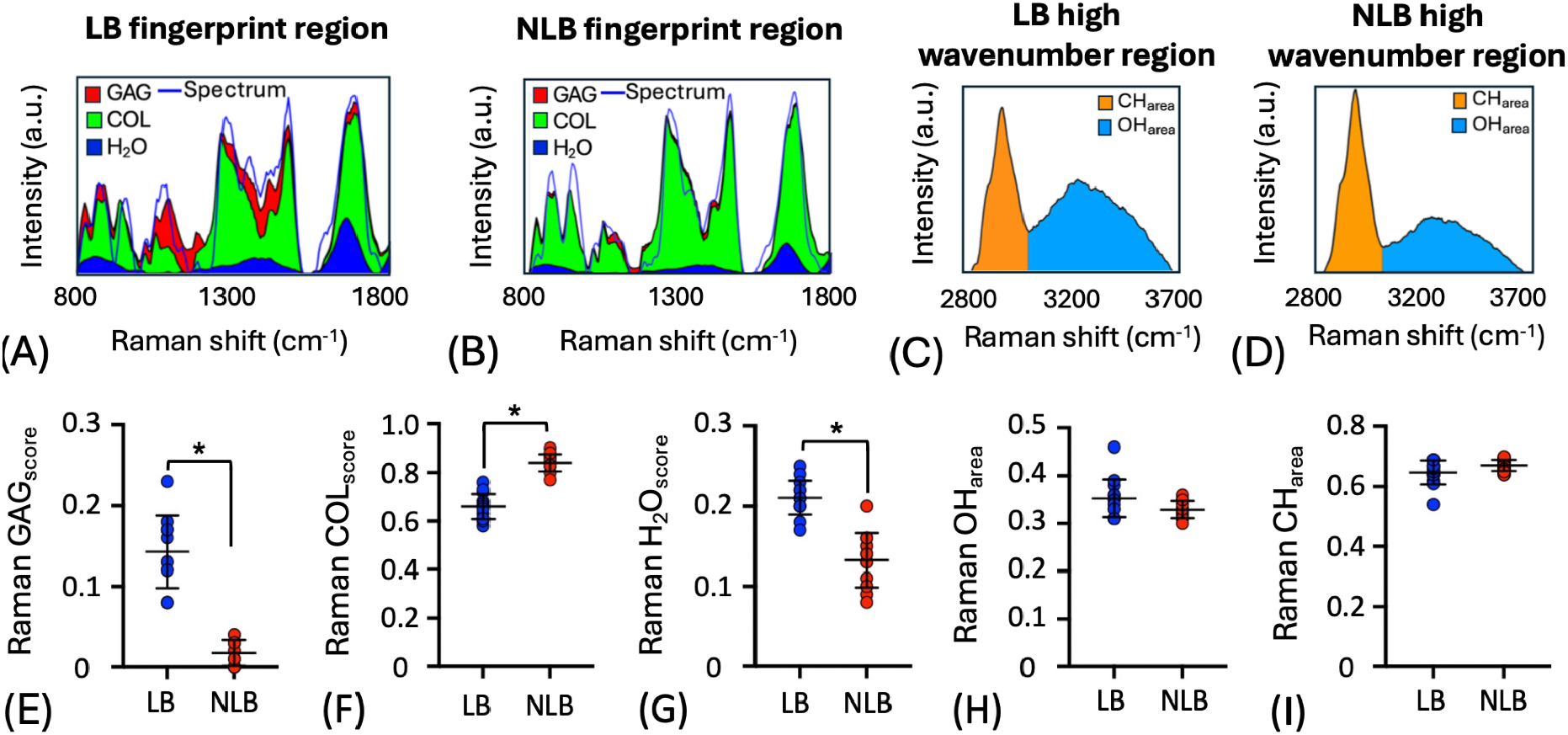
Raman spectral analysis of LB and NLB equine osteochondral explants. **(A, B)**. Raman spectra derived biomarkers representing cartilage ECM content of the uppermost 300 μm of the chondral layer portraying the superficial and middle zones. Representative stacked area plots indicating the relative contribution of GAG, COL, and water to the composite cartilage fingerprint Raman spectra of LB medial condyle and NLB trochlear groove explants. **(C, D)** Representative high wavenumber Raman spectra, depicting spectral CH_area_ and OH_area_ of LB and NLB cartilage. Comparative analysis of Raman biomarkers derived from fingerprint spectral analysis for LB vs NLB cartilage: **(E)** Raman GAG_score_, **(F)** Raman COL_score_, **(G)** Raman H_2_O_score_. **(H)** Raman CH_area_ and **(I)** Raman OH_area_. LB cartilage ECM demonstrated significantly higher GAG content, significantly lower COL content, and significantly higher water content compared to NLB cartilage ECM (*p* < 0.05).

### Raman correlations with cartilage properties

Raman-spectra-derived biomarker scores accounted for changes in the composition and structure of cartilage explants (**Table 1**). GAG_score_ accounted for 81% of the variation of digital-densitometry-measured GAG content (**Fig.5A**; R^2^=0.81; *p* < 0.001). COL_score_ accounted for 53% of the variation in FTIR-measured collagen content (**Fig.5B**; R^2^=0.53; *p* < 0.001) and 59% of the variation in collagen parallelism index (**Fig.5C**; R^2^=0.59; *p* < 0.001). Biomarker scores accounted for changes in the viscoelastic properties of cartilage explants. GAG_score_ accounted for 78% of the variation equilibrium modulus (**Table 1**; R^2^=0.78; *p* < 0.001) and 44% of the variation in phase shift angle (**Table** 1; R^2^=0.44; *p* < 0.001). A linear combination of GAG_score_ and OH_area_ accounted for 81% variation of equilibrium modulus (**Fig.5D**; R^2^=0.81; *p* < 0.001), 35% variation of dynamic modulus (**Table 1**; R^2^=0.35; *p* < 0.01), and 51% of the variation in phase shift angle (**Fig.5F**; R^2^=0.51; *p* < 0.001). A linear combination of GAG_score_ and COL_score_ accounted for 58% variation of dynamic modulus (**Fig.5E**; R^2^=0.58; *p* < 0.001). Raman biomarker scores accounted for variation in the cartilage biphasic properties. The linear combination of GAG_score_ and OH_area_ accounted for 83% of the variation in the non-fibrillar matrix modulus (**Fig.5G**; R^2^=0.83; *p* < 0.001). The linear combination of GAG_score_ and COL_score_ accounted for 19% of the variation in the fibril network modulus (**Fig.5H**; R^2^=0.19; *p* < 0.05) and 34% of the variation in Darcy permeability (**Fig.5I**; R^2^=0.34; *p* < 0.01).

**Table 1:** Coefficients of determination for univariate and multivariate correlations between Raman biomarker scores and composition (GAG content, COL content), structure (parallelism index [*PI*]), viscoelastic properties (equilibrium modulus [*E*_*eq*_], dynamic modulus [*E*_*dyn*_], phase shift [*θ*], and biphasic properties (non-fibrillar matrix modulus [*E*_*nf*_], Darcy permeability [*k*_*0*_], fibrillar network modulus [*E*_*f*_]) of equine explants. Coefficients of determination that explain a significant amount of variation (p<0.05) indicated in bold.

| **Tissue property** |  | **Univariate correlations** |  |  |  |  | **Multivariate correlations** |  |  |
|---|---|---|---|---|---|---|---|---|---|
|  |  | **Raman Biomarker** |  |  |  |  | **Raman Biomarker** |  |  |
|  |  | **Fingerprint score** |  |  | **High wavenumber area** |  | GAG<sub>score</sub> + OH<sub>area</sub> | COL<sub>score</sub> + OH<sub>area</sub> | GAG<sub>score</sub> + COL<sub>score</sub> |
|  |  | GAG | COL | H<sub>2</sub>O | OH<sub>area</sub> | CH<sub>area</sub> |  |  |  |
| **Composition & structural properties** | ***GAG*** | **0.81<br>p<0.001** | **0.77<br>p<0.001** | **0.54<br>p<0.001** | 0.05<br>p=0.30 | 0.06<br>p=0.25 | **0.86<br>p<0.001** | **0.77<br>p<0.001** | **0.87<br>p<0.001** |
|  | ***COL*** | **0.60<br>p<0.001** | **0.53<br>p<0.001** | **0.43<br>p<0.001** | 0.08<br>p=0.18 | 0.09<br>p=0.15 | **0.65<br>p<0.001** | **0.52<br>p<0.001** | **0.58<br>p<0.001** |
|  | ***PI*** | **0.44<br>p<0.001** | **0.59<br>p<0.001** | **0.50<br>p<0.001** | 0.05<br>p=0.30 | 0.06<br>p=0.25 | **0.45<br>p<0.001** | **0.53<br>p<0.001** | **0.53<br>p<0.001** |
| **Viscoelastic properties** | ***E<sub>eq</sub>*** | **0.78<br>p<0.001** | **0.72<br>p<0.001** | **0.31<br>p<0.01** | **0.21<br>p<0.05** | **0.18<br>p<0.05** | **0.81<br>p<0.001** | **0.73<br>p<0.001** | **0.81<br>p<0.001** |
|  | ***E<sub>dyn</sub>*** | **0.54<br>p<0.001** | **0.34<br>p<0.01** | 0.01<br>p=0.64 | **0.30<br>p<0.01** | 0.13<br>p=0.08 | **0.35<br>p<0.001** | **0.36<br>p<0.01** | **0.58<br>p<0.001** |
|  | ***θ*** | **0.44<br>p<0.001** | **0.42<br>p<0.001** | **0.48<br>p<0.001** | 0.11<br>p=0.11 | 0.13<br>p=0.08 | **0.51<br>p<0.001** | **0.43<br>p<0.001** | **0.44<br>p<0.001** |
| **Poroelastic properties** | ***E<sub>nf</sub>*** | **0.83<br>p<0.001** | **0.73<br>p<0.001** | **0.34<br>p<0.01** | **0.18<br>p<0.05** | **0.18<br>p<0.05** | **0.83<br>p<0.001** | **0.74<br>p<0.001** | **0.82<br>p<0.001** |
|  | ***k<sub>0</sub>*** | **0.26<br>p<0.01** | **0.17<br>p<0.05** | 0.06<br>p=0.25 | 0.11<br>p=0.11 | 0.01<br>p=0.64 | **0.29<br>p<0.01** | **0.20<br>p<0.05** | **0.34<br>p<0.01** |
|  | ***E<sub>f</sub><sup>0</sup>*** | 0.12<br>p=0.10 | 0.06<br>p=0.25 | 0.01<br>p=0.64 | **0.25<br>p<0.05** | **0.25<br>p<0.05** | 0.13<br>p=0.08 | 0.08<br>p=0.18 | **0.19<br>p<0.05** |

## Discussion

Raman-probe-derived biomarkers for GAG, COL and water accurately portrayed the major compositional constituents of the cartilage ECM and account for the viscoelastic and poroelastic material properties fundamental to articular cartilage function. To establish the sensitivity of Raman biomarkers to describe subtle compositional and structural variations, we analyzed explants from high load bearing (medial femoral condyle) and low load bearing (intertrochlear groove) anatomic regions. Consistent with its elevated load-bearing mechanical demands, the LB cartilage was stiffer and more elastic (lower phase shift angle, *p* < 0.05) than the NLB cartilage. These functional differences are attributed to a higher tissue GAG content in LB cartilage, which generates elevated Donnan osmotic swelling pressure, complemented by a trend of lower permeability (*p* = 0.06) that restricts fluid flow and further augments load support during dynamic compression.

ECM-specific Raman biomarkers accurately depicted these site-dependent compositional variations. GAG_score_ accounted for 81% of the variation in GAG content, and COL_score_ accounted for 53% of the variation in collagen content. Interestingly the COL_score_ additionally accounted for 59% of the variation of the collagen parallelism index, reflecting collagen fibril anisotropy, a structural feature known to deteriorate with collagen fibrillation in early-stage OA.^59^ This correlation likely reflects a loss of superficial zone collagen alignment at the high load bearing medial femoral condyle compared to the trochlear groove cartilage (**Fig. 5C)**. Future polarized Raman analyses will measure collagen birefringence to improve anisotropy quantification.^41^ Water-related biomarkers (Raman H_2_O_score_ and high wavenumber OH_area_) were lower in NLB cartilage, consistent with site-dependent biochemical variation. While tissue porosity was not directly measured in this study, these parameters correlated with water content in prior work.^40^

**Figure 5.**
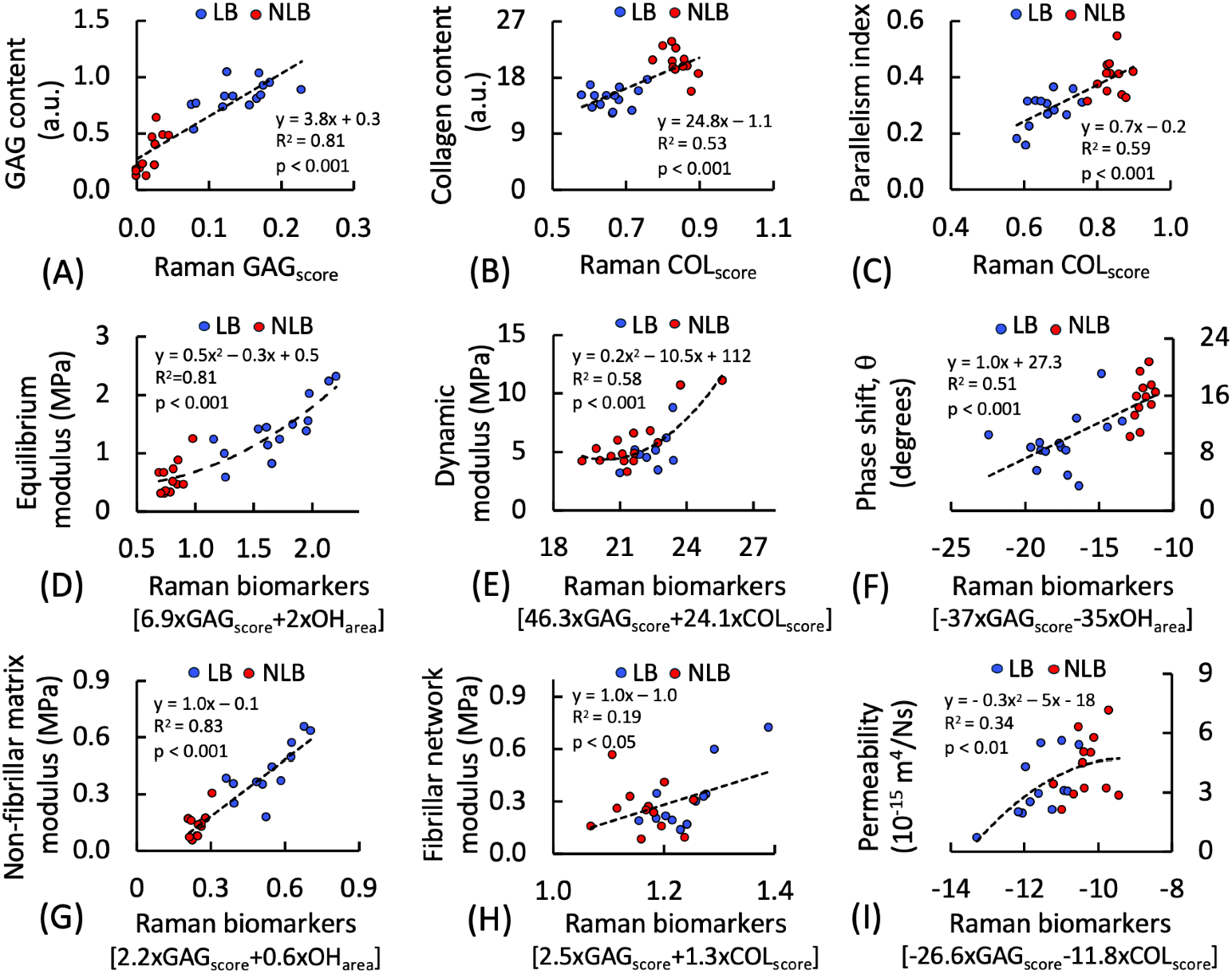
Bivariate correlation between Raman biomarker scores and cartilage composition, structure, viscoelastic, and biphasic properties of LB and NLB equine osteochondral explants.

Raman ECM biomarkers representing the GAG matrix and collagen fibril network account for the viscoelastic material properties fundamental to cartilage load-bearing function. A linear combination of GAG_score_ and OH_area_ accounted for 81% of the variation in equilibrium modulus and 51% of the variation in phase shift, a measure of viscoelastic energy dissipation during dynamic loading. These associations are consistent with the established role of GAG-associated Donnan osmotic swelling pressure in resisting compressive deformation and promoting IFLS. A linear combination of GAG_score_ and COL_score_ accounted for 58% of the variation in dynamic modulus, reflecting the combined contributions of proteoglycan-associated fluid pressurization and collagen network reinforcement, which together regulate cartilage load support under dynamic compression.^7;46;52^ These relationships are mechanistically supported by the coupled roles of GAG and the collagen network in regulating cartilage mechanical behavior. GAG depletion reduces Donnan osmotic swelling pressure, compromises IFLS, diminishes electrostatic repulsion among anionic sulfate groups,^60^ and increases tissue porosity and interstitial fluid content. Collectively, these changes can impair resistance to rapid compressive deformation and render cartilage more vulnerable to degeneration under repetitive, high-frequency loading, such as running.^46;52^ Thus, Raman-derived biomarkers provide an objective means of characterizing cartilage mechanical competency from its ECM composition and structure.

We employed a fibril-reinforced biphasic FE model to evaluate the contributions of specific ECM constituents to the solid-fluid interactions governing cartilage’s poroelastic behavior. A linear combination of GAG_score_ and OH_area_ accounted for 83% of the variation in the non-fibrillar matrix modulus, demonstrating a strong association between Raman-derived biomarkers and GAG-associated material properties. The lower GAG content detected at the trochlear groove NLB cartilage was associated with a significant reduction in the GAG-associated non-fibrillar matrix modulus and a trend toward increased Darcy permeability (*p* = 0.06). These findings mechanistically align with the established relationship between GAG content, non-fibrillar matrix modulus, and fluid transport. No significant difference was observed in the fibrillar network modulus between cartilage sites. Fibrillar modulus estimates may be limited by the sensitivity of the FE model, particularly if interstitial fluid pressurization was inadequate to engage the collagen network sufficiently to reveal changes in fibrillar stiffness. Also, polarized light microscopy confirmed that collagen anisotropy was lower in the medial condyle LB cartilage than the trochlear groove NLB cartilage, which can influence macroscopic material behavior (e.g., dynamic modulus), but was not captured by the fibril-reinforced biphasic FE model which assumed an idealized uniformly aligned fibrillar architecture and did not incorporate sample-specific variations in collagen orientation and anisotropy. As such, the derived fibril network modulus represents only the intrinsic stiffness of the collagen fibrils. In the future, additional Raman analysis of the collagen microstructure, including crosslinks, collagen alpha helix stability,^29^ and polarized Raman measures of collagen alignment,^41^ may account for a greater part of the variation in collagen fibril network properties.

A limitation of this study is that the experiments were conducted using equine cartilage samples. Although equine articular cartilage shares many structural and compositional similarities with human cartilage, the findings cannot be directly extrapolated to human tissue. Species-specific differences in cartilage composition and architecture within the superficial cartilage layer may influence the correlation estimates. The use of macroscopically normal cartilage with subtle compositional and structural variations across anatomical sites enabled rigorous validation of biomarker sensitivity to physiologically relevant compositional changes in the early stages of cartilage degeneration. However, it did not account for more severe degeneration associated with chondral lesions. Consequently, the limited dynamic range of tissue properties may underestimate the association of biomarkers in pathologically degenerated cartilage, where GAG loss and collagen network disruption are more pronounced. A second limitation stems from the probe’s interrogation depth, as this analysis examined only the superficial and middle cartilage zones (topmost 300μm). While this region is sensitive to early-stage degeneration, it provides limited information about deeper structural changes. In future work, deeper-penetrating lenses can be adopted to achieve simultaneous compositional analysis of deeper tissue regions, as we have performed previously.^40^ A third limitation arises from spatial heterogeneity in the correlative analysis. Although we aligned regions of analysis between Raman and compositional measures by averaging values over the probe interrogation window (topmost 300μm), this comparison remains imperfect due to the nonuniformity of the Raman interrogation region and the nonlinear, depth-dependent variation in tissue composition, which could affect correlation estimates.

The results of this study support the translation of Raman spectroscopy as a clinically transformative diagnostic platform for objective cartilage assessment during surgical repair procedures. The arthroscopy-compatible Raman needle probe can be utilized inter-operatively to evaluate cartilage tissue, enabling: 1) objective delineation of diseased cartilage margins based on compositional biomarkers that mechanistically account for biomechanical competency, and 2) real time quantification of repair tissue composition during second-look arthroscopy to objectively evaluate repair tissue and guide retreatment decisions. By providing standardized quantitative biomarkers that mechanistically reflect tissue function rather than subjective surface assessment, Raman spectroscopy enables objective diagnosis of cartilage injury, interoperative monitoring during repair, and evaluation of therapeutic interventions.

Raman biomarkers further offer advantages as standardized, objective measures of cartilage health across the evolving hierarchy of tissue models, from *ex vivo* explant studies to preclinical *in vivo* animal studies to clinical trials. For *ex vivo* studies, the platform’s low cost, portability, and ease of use make Raman probes attractive alternatives to laborious biochemical assays and histological staining, enabling rapid, non-destructive compositional assessment. Critically, non-destructive Raman measurement permits serial monitoring of live cartilage explants in culture in response to mechanical and chemical insults and drug therapies. For *in vivo* preclinical studies, Raman probes enable the minimally invasive arthroscopic assessment of cartilage composition in large animals, permitting longitudinal monitoring of disease progression and therapeutic response. For clinical trials, Raman-derived biomarkers can serve as the objective primary endpoint measure to quantify the efficacy of therapeutic interventions for inhibiting cartilage degeneration or promoting repair.

The demonstrated link between Raman-derived compositional biomarkers and functional tissue properties establishes a mechanistic foundation for Raman spectroscopy to advance cartilage diagnostics. By standardizing cartilage composition assessment through non-destructive, objective biomarkers that mechanistically account for tissue function, Raman spectroscopy has the potential to accelerate the development and clinical translation of novel therapeutic strategies for cartilage preservation and repair.

## Acknowledgements

Research reported in this publication was supported by the National Institute of Arthritis and Musculoskeletal and Skin Diseases under award number R01AR081393, the Arthritis Foundation, MTF Biologics, the Boston University Materials Science & Engineering Innovation Award, Maud Kuistila muistosäätiö (0234A2), State Research Funding for university-level health research, Kuopio University Hospital, Wellbeing Services County of North Savo (5063599), Research Council of Finland (348410, 357787, 372127), Regional Council of Pohjois-Savo (A74798), and Biocenter Kuopio. The opinions, findings, and conclusions, or recommendations expressed are those of the authors and do not necessarily reflect the views of the National Institutes of Health.

## Author Contributions

**Massie Kazemi**: conceptualization, data acquisition and analysis, validation, writing – original draft. **Juuso Tuppurainen**: conceptualization, data acquisition and analysis, validation, writing – original draft. **Jiri Jäntti:** data acquisition and analysis, writing – review and editing. **Chenhao Yu:** data acquisition and analysis, writing – review and editing **Maria C. Fugazzola:** sample preparation, data acquisition and analysis, writing – review and editing. **René V. Weeren:** research design, resources, writing – review and editing. **Thomas P. Schaer:** research design, writing – review and editing. **Mark W. Grinstaff:** research design, writing – review and editing. **Mads S. Bergholt:** conceptualization, writing – review and editing. **Brian D. Snyder:** conceptualization, research design, supervision, writing – original draft. **Janne T.A. Makela:** conceptualization, research design, resources, supervision, writing – original draft, funding acquisition. **Michael B. Albro:** conceptualization, research design, resources, supervision, writing – original draft, funding acquisition. All authors read and approved the final manuscript.

